# Spritz: A Proteogenomic Database Engine

**DOI:** 10.1101/2020.06.08.140681

**Authors:** Anthony J. Cesnik, Rachel M. Miller, Khairina Ibrahim, Lei Lu, Robert J. Millikin, Michael R. Shortreed, Brian L. Frey, Lloyd M. Smith

**Affiliations:** Department of Chemistry, University of Wisconsin-Madison, Madison, WI, USA; Science for Life Laboratory, School of Engineering Sciences in Chemistry, Biotechnology and Health, KTH - Royal Institute of Technology, Stockholm, 17121, Sweden; Department of Genetics, Stanford University, Stanford, CA 94305, USA; Chan Zuckerberg Biohub, San Francisco, CA 94158, USA

**Keywords:** RNA-Seq, proteogenomics, sequence variations, modifications, proteoform, top-down, sample-specific, transcriptomics, PTMs

## Abstract

Proteoforms are the workhorses of the cell, and subtle differences between their amino acid sequence or post-translational modifications (PTMs) can change their biological function. To most effectively identify and quantify proteoforms in genetically diverse samples by mass spectrometry (MS), it is advantageous to search the MS data against a sample-specific protein database that is tailored to the sample being analyzed, in that it contains the correct amino acid sequences and relevant PTMs for that sample. To this end, we have developed Spritz (https://smith-chem-wisc.github.io/Spritz/), an open-source software tool for generating protein databases annotated with sequence variations and PTMs. We provide a simple graphical user interface (GUI) for Windows and scripts that can be run on any operating system. Spritz automatically sets up and executes approximately 20 tools, which enable construction of a proteogenomic database from only raw RNA sequencing data. Sequence variations that are discovered in RNA sequencing data upon comparison to the Ensembl reference genome are annotated on proteins in these databases, and PTM annotations are transferred from UniProt. Modifications can also be discovered and added to the database using bottom-up mass spectrometry data and global PTM discovery in MetaMorpheus. We demonstrate that such sample-specific databases allow the identification of variant peptides, modified variant peptides, and variant proteoforms by searching bottom-up and top-down proteomic data from the Jurkat human T lymphocyte cell line and demonstrate the identification of phosphorylated variant sites with phosphoproteomic data from the U2OS human osteosarcoma cell line.

## Introduction

Proteins comprise the machinery of the cell, performing a multitude of functions. These molecules can be modified by a host of enzymes such as kinases, creating a variety of proteoforms with unique activities and functions. To understand the functions of proteoforms^1^ and their roles in the cell, we must identify and quantify them. Top-down mass spectrometry (MS) is a proteomic method that can directly analyze proteoforms ^2,3^, and bottom-up MS analysis can approximate this type of study by analyzing peptides from these proteoforms. We previously demonstrated how using a database annotated with modifications discovered in bottom-up proteomic analysis expands the number of proteoform identifications ^4,5^. In this work, we demonstrate how creating databases with full length proteoform sequences containing variations can improve the identification of amino acid sequence variations, allow identification of modified variant peptides, and enable the identification of variant proteoforms.

The rising tide of computational workflows that aim to detect amino acid sequence variations by leveraging nucleotide sequencing data has collectively been termed proteogenomics ^6,7^. There are several barriers for the wide-spread adoption of proteogenomic methods in the proteomics community including the steep learning curve for genomics tools and the difference in operating system requirements for genomics and proteomics tools. Typically, tools for proteomics are written for Windows using languages like C#. Tools for genomics are written to run in Unix command-line interfaces (*e.g.,* Linux, Mac). This conflict leads to a difficult problem of integrating these tools. MaxQuant has solved this problem by using .NET Core ^8^ to allow the program to run in Unix environments, and tools like ProteomeGenerator ^9^ have used Docker or Singularity to place the program in a container that can run on any system. In this work, we focus on making these tools easy for the proteomics community to run in Windows environments, which are more common among proteomics labs. In doing so, we developed Spritz, which is written in the programming language *snakemake*. Snakemake facilitates the installation and dependency management for these tools and simplifies the writing of scripts to run these tools given any input files or publicly available data ^10^. We also created a graphical user interface (GUI) for Windows to ease the learning curve by packaging the *snakemake* workflows, which can be challenging to edit, into a point-and-click type program to create proteogenomic databases.

Tools for creating proteogenomic databases with sequence variations typically require preprocessed data, either in variant call format (VCF) files or binary alignment map (BAM) files, such as in the variant extractor tool within MaxQuant ^8^. However, there are many steps to perform prior to getting these files. In Figure 1, we show our workflow for analyzing genomics data. Spritz facilitates this entire analysis from raw input to output.

**Figure 1:**
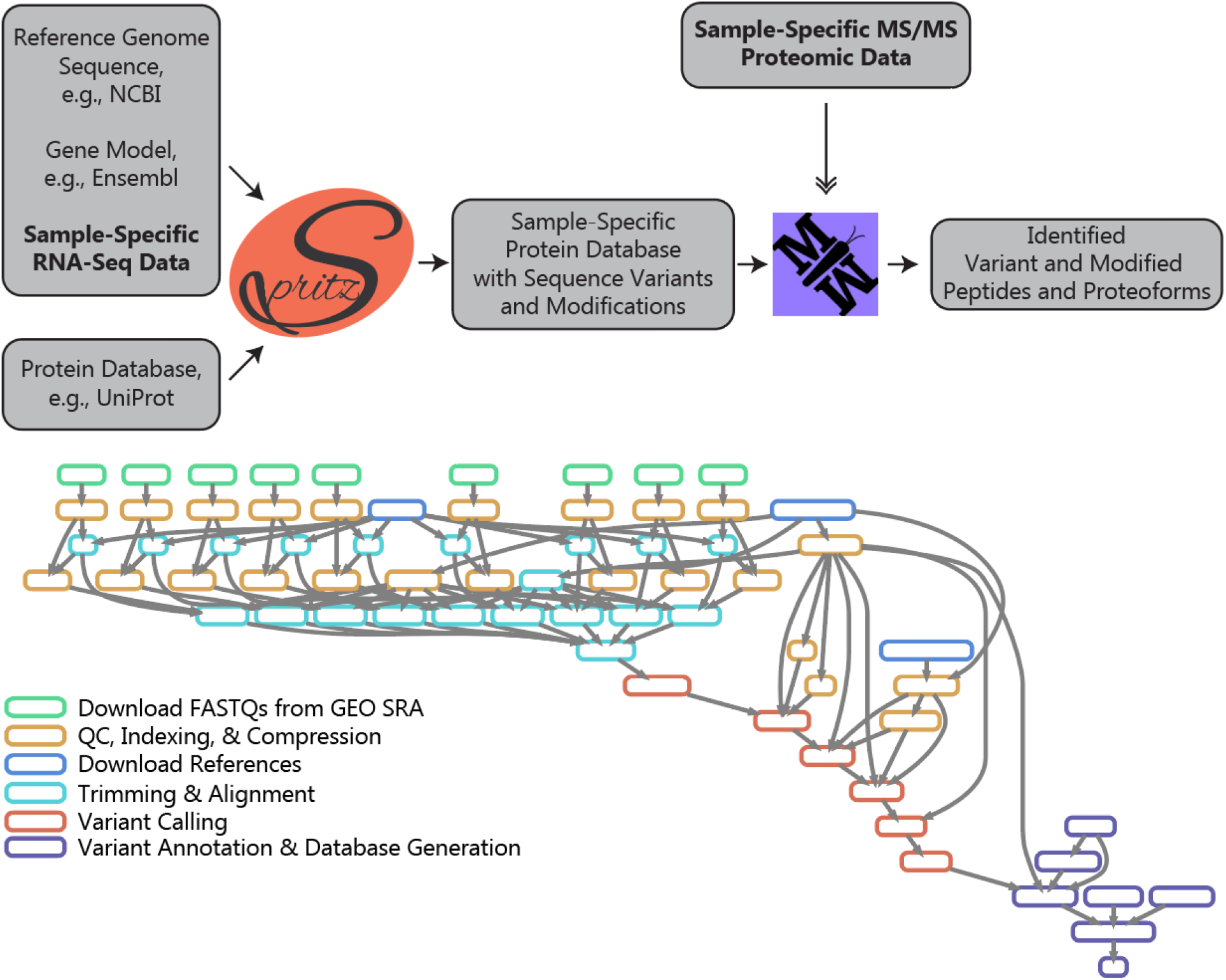
(Two columns.) Spritz is a pipeline of approximately 20 tools that produces sample-specific protein databases annotated with modifications and sequence variations. Spritz automatically downloads the genome fasta file, gene model file, and UniProt XML protein database for the organism. If publicly available sequence read archives (SRAs) are available for the sample of interest, Spritz will also download those raw RNA-Seq data. MetaMorpheus is used in this work to search both bottom-up and top-down mass spectrometry proteomics data. Bottom-up proteomics data is used to discover additional sites of modification to help enable the identification of variant proteoform sequences.

Historically, identifying variant peptides was performed by parsing variant calls into peptides (*e.g.,* tryptic peptides) and then appending those peptides to the database ^11–15^. This approach precludes the identification of full length proteoforms, and so in this work, we apply variations to the entire protein sequence. By entering variants into the entire protein sequence one at a time, we can replace the reference allele completely for homozygous variations and generate combinations of heterozygous variations. This reduces false positive identifications of the reference allele for homozygous variations when compared to strategies that append the variant peptide to the database without removing the reference allele. It may also enable the detection of proteoforms from different alleles of the same gene by including combinations of heterozygous variations.

Proteogenomic workflows typically only look for single amino acid variations (SAAVs) by filtering for missense single nucleotide variations (SNVs). This ignores the complex landscape of indels and frameshift peptides, thereby precluding the investigation of the effect these common changes have on protein functions. Spritz removes this restriction and enables the analysis of peptides resulting from multiple nucleotide variants (MNVs), small indels, frameshifts and other types of variations. Calling MNVs is an important and sometimes overlooked step in appending missense mutations for proteomic analysis, since SNVs that appear within 3 bases of one another can change a codon together.

Finally, proteomic searches have recently become increasingly capable of discovering sites of modification and identifying modified peptides ^16–19^, and they are thus also capable of identifying modified variant peptides given a sample-specific database. However, no other search strategies, to our knowledge, are capable of detecting post-translationally modified variant sites. In this work, we adapted the protein sequence database format for UniProt XMLs to enable the annotation of modifications on amino acids within a variant site.

## Materials and Methods

Spritz is an open-source pipeline written in *snakemake* that facilitates proteogenomic data analysis by taking in FASTQ files from RNA-Seq analysis or publicly available SRA accessions (Figure 1; https://smith-chem-wisc.github.io/Spritz/). Spritz uses an adapted version of the UniProt-XML format that can hold variant annotations, site-specific modification annotations, and modification annotations on variants. This was implemented in mzLib (a library for MS data analysis, located at https://github.com/smith-chem-wisc/mzLib) and MetaMorpheus^20^ to enable searching for modified and variant peptides, proteoforms, and annotating new sites of modifications on proteins and variants.

### Data Sets

Jurkat data were obtained previously: RNA-Seq data (GSE45428 in GEO SRA)^12^ consisted of paired-end reads 101 nucleotides in length, and bottom-up proteomics data consisted of 28 fractions of high-pH reverse phase liquid chromatography (RPLC) from a tryptic digestion of Jurkat cell lysate as well as 11 fractions for five other proteases^12^. The second dataset used to assess the identification of modified variant sites is a phosphoproteomic analysis of U2OS cells^21^, along with U2OS RNA-Seq data obtained from the Human Protein Atlas ^22^ (SRR629563 in GEO SRA). Finally, a recently published top-down proteomics dataset for Jurkat was used to demonstrate the identification of variant-containing proteoforms ^5^.

### Installing Genomics Tools and Dependencies

Spritz facilitates the installation of genomics tools using Bioconda ^23^ by creating and activating an environment for the tools within a Unix command-line environment. The tools are also packaged into a Docker container using Conda, which is downloaded and executed by the GUI.

### Downloading References and Sequence Read Archives

Spritz automatically downloads genomic references needed for genomic analysis, including the human genome (FASTA format) and gene model (GFF3 format) files from Ensembl version 81. Known sites of variation in humans are downloaded from dbSNP ^24^ (version human_9606_b150_GRCh38p7). Paired-end FASTQ files from GEO SRA are downloaded using the tool *fasterq-dump* (Table S1) ^25^.

### Preprocessing FASTQ files and Quality Control

Spritz uses the program *skewer* ^26^ to trim adapter sequences from the RNA sequences using a list of adapters obtained from BBMap. *Skewer* also filters low quality reads, and in this work, we require an average base quality score of Q=20 for each read. Spritz uses the program *fastqc* to get quality metrics for each file before and after trimming the reads.

### Aligning RNA-Seq Reads

Spritz uses *hisat2* ^27^ to align reads to the human reference genome (Ensembl GRCh38 version 81 in this work). We chose this software because of its low RAM requirements, since typical machines used for proteomics have around 16 GB of RAM. In this work, we analyzed the FASTQ files from each SRA accession separately, and then combined the resulting BAM files for variant calling analysis.

### Variant Calling

The aligned reads are prepared for variant calling analysis using the Genome Analysis Toolkit, GATK ^28–30^ (version 4.0.11.0). The reads in the combined BAM file are grouped into a single group using ‘gatk AddOrReplaceReadGroups,’ and then the duplicate reads are marked using ‘gatk MarkDuplicates.’ After detecting whether base quality scores are consistent with updated standards using ‘gatk FixMisencodedBaseQualityReads,’ the RNA-Seq reads that are aligned across splice junctions are split with a tool named ‘gatk SplitNCigarReads.’ Base quality scores are then recalibrated using known sites of variation using ‘gatk BaseRecalibrator’ and ‘gatk ApplyBQSR.’

After these steps of preparation, variants are called at each site in the genome using ‘gatk HaplotypeCaller’ to produce a genome variant call file (gVCF). A minimum base quality score of 20 is required, and soft clipped bases are not used in this analysis. Multi-nucleotide variants (MNVs) are called using the setting ‘--max-mnp-distance 3,’ which was chosen to match the length of a codon to capture all possible codon changes. The gVCF is indexed with ‘gatk IndexFeatureFile,’ and then the file is genotyped for Jurkat or U2OS using ‘gatk GenotypeGVCFs’ to produce a raw VCF file.

### Variant Annotation and Proteogenomic Database Construction

Designing software to analyze the effects of protein coding variations requires an intricate data structure representing the genome and gene model. The variant annotation software SnpEff ^31^ has code to represent these structures, including codons, spliced transcripts, full length mRNAs including UTRs, genes, and chromosomes. In this work, we created an adapted version of SnpEff (located at https://github.com/smith-chem-wisc/SnpEff) to annotate variants within a protein database. To do this, we adapted the UniProt-XML format to enable protein sequence variant annotations. These variant features contain the original amino acid sequence, the variant amino acid sequence obtained from applying the variation within SnpEff, and the entire line from the VCF file from which it originated. These annotations are used within MetaMorpheus to create variant protein sequences. SnpEff uses the raw VCF as input and outputs a VCF file with annotations from Ensembl (version 86) and a UniProt-XML database containing proteins translated from Ensembl transcripts annotated with coding variations, such as missense mutations and indels.

Spritz then automatically downloads a reference UniProt-XML database from UniProt ^32^ (downloaded April 18, 2019 for Jurkat and May 13, 2020 for U2OS) and annotates modifications, gene names, and other information for these protein sequences that match UniProt exactly using a module named TransferUniProtModifications. This module is written in C# to take advantage of the data structure representations of proteins and modifications within mzLib.

### Augmented UniProt-XML Format for Modified Sequence Variant Annotation

The G-PTM-D strategy was amended in MetaMorpheus and the underlying mzLib software library was updated to output a modified UniProt-XML protein database format that allows the annotation of modifications that are discovered on sites of sequence variations. Specifically, when a candidate modification site overlaps with an applied variation, a modification ‘subfeature’ is added to the ‘location’ annotation for the sequence variation ‘feature.’ This allows MetaMorpheus to generate these modified variant forms within the search space for both bottom-up and top-down proteomic searches.

The stability of these databases is also improved over the UniProt-XML format by including the properties of each modification within the database rather than referencing separate PTM lists that are subject to change (*e.g.,* https://www.uniprot.org/docs/ptmlist).

### Bottom-Up Mass Spectrometry Analysis

MetaMorpheus (version 303 for Jurkat and 305 for U2OS) was used to analyze bottom-up proteomic data generated with multiple proteases including 28 high-pH fractions^12^ for trypsin and 11 fractions for each of LysC, AspN, ArgC, GluC, and chymotrypsin ^11^. These data were analyzed with the Spritz-generated proteogenomic database constructed with variations applied to the Ensembl reference. First, MetaMorpheus was used to calibrate RAW files using the settings listed in Table S2, producing mzML files.

Then, the calibrated files were analyzed with the global post-translational modification discovery (G-PTM-D) search within MetaMorpheus ^17,20^ (settings listed in Table S2) to identify and annotate candidate modification sites for common biological modifications (*e.g.,* acetylation, phosphorylation), metal adducts (e.g. sodium, iron), and sample preparation artifacts (*e.g.,* ammonia loss, deamidation).

G-PTM-D produces a new database containing annotations for the discovered candidate PTMs. This updated proteogenomic database was used to search the calibrated spectra files using tighter mass tolerances calculated by MetaMorpheus after calibration (5.8 ± 2.6 PPM precursor mass tolerance and 10.7 ± 1.9 PPM product mass tolerance, improved from ± 15 PPM and ± 25 PPM before calibration, respectively; Table S2). The results of this search include peptide spectral match (PSM), peptide, and variant peptide files. To identify a sequence variant, we required peptides to identify only variant proteins, *i.e.,* not also be a potential digestion product of a non-variant or canonical protein. We also required the peptide sequence to intersect the variant or to identify a new proteolytic cleavage site as a result of the variation. The search task also produces a “pruned” database and a “protein pruned” database. The “pruned” database contains only modifications identified by peptides at 1% FDR, and the “protein pruned” database only contains protein sequences and modifications that were identified by peptides at 1% FDR. The simplified “pruned” database was used for top-down proteoform analysis in this work.

### Top-Down Mass Spectrometry Analysis

MetaMorpheus was also used for top-down proteomic analysis. The “pruned” database containing only modifications detected at a 1% FDR level was used to search the 11 GeLFREE ^33^ fractions using the settings listed in Table S2. The results of this search are lists of the proteoform spectral matches (PrSM) and proteoforms, as well as the variant-containing proteoforms listed in accompanying files.

## Results

### Ease of Use

Spritz eases the learning curve of RNA-Seq proteogenomics in several ways: 1) it allows raw RNA-Seq reads to be used as input rather than downstream results files; 2) it automates the execution of approximately 20 genomic analysis tools; 3) it improves the compatibility between tools for genomic and proteomic analysis; and 4) it provides a GUI for Windows that eliminates the need for users in the proteomic community to operate command-line interfaces. While the raw material for proteogenomics is nucleotide sequencing data, tools for proteogenomic analysis typically require pre-processed files. As described in the Methods section, producing these files is not trivial and requires a pipeline of over approximately 20 tools for preprocessing, aligning, and variant calling. Spritz solves this problem by using the programming language *snakemake* to take raw RNA-Seq reads as input and automatically run these tools before using custom Spritz code to generate proteogenomic databases. A similar approach using *snakemake* was used by the proteogenomic program ProteomeGenerator ^9^ that focuses on novel isoform detection. However, that program does not provide a user interface, which raises the challenge of reusing the pipeline by requiring users to edit *snakemake* files.

The second barrier for entry in using proteogenomic databases in proteomics is a mismatch between the operating systems required for various tools enabling genomic and proteomic analyses. Typically, genomic tools are run in Unix environments, such as Macintosh and Linux, whereas proteomic tools typically are built for Windows. While solutions exist for bridging this gap, such as Docker and Singularity, using these “containerizing” programs is not trivial for beginning users and adds difficulty to the challenge of learning how to combine results from various genomic tools.

Most tools for genomic analysis use the command-line interface for executing the program. The command-line interface is an excellent tool for developers and data analysis experts for passing settings or files to the program with only a few lines of code. However, the command-line interface can be intimidating and difficult for beginners to use on its own, let alone in addition to the learning curve for using each program. Existing solutions to remedy this problem include Galaxy-P^13,34^, which has been used previously to string together tools for proteogenomics. We chose to solve this problem using a GUI written in C# that runs in Windows and operates a Docker container containing the *snakemake* scripts. The GUI also allows the user to select the organism and gene model version, so that Spritz can automatically download the proper Ensembl and UniProt reference files. Altogether, this means that the user will need to download Docker and Spritz but will not necessarily need to know how to run the Docker container or edit *snakemake* scripts on their own, making proteogenomic analysis more accessible to the proteomics community.

In this work, we demonstrate the use of Spritz to find variant peptides, modified variant peptides, and variant proteoforms in Jurkat cell lysate, as well as phosphorylated variant peptides in a phosphoproteomic dataset for U2OS cell lysate, from previously published bottom-up and top-down results.

### Variant Calling and Proteogenomic Database Creation

The pre-processing and alignment of the RNA-Seq reads allows us to use tools for calling variants against the reference genome. The entire Jurkat RNA-Seq dataset contained 113,399,653 reads. After trimming and filtering, 110,663,075 reads remained, and the mean per base quality score increased from 16.3 to 17.3, and 95.3% of these reads were aligned to the genome by *hisat2*.

Variant calling was performed by GATK HaplotypeCaller, which found 371,784 variants (one variant every 8,326 bases of the genome). A customized version of SnpEff was used to generate 60,007 protein sequences from the Ensembl reference genome, which were annotated with 22,930 coding variations (Figure 2). Among the most common single amino acid variations (SAAVs) were R > H, R > Q, and R> W, which eliminate tryptic cleavage sites. The 60,007 protein sequences and 22,930 coding variations were used to generate a search space within MetaMorpheus containing 71,328 variant proteins with 22,407 of these variations (Figure 3A). In this process, we filtered the 496 variations that do not lead to meaningful sequences, such as frameshifts that truncate the protein sequence to below 7 amino acids in length. A total of 49,999 modified residue annotations (95% of 52,643) of 114 modification types were transferred from a UniProt-XML database for the 73% of proteins (43,636/60,007) with exact sequence matches in UniProt. These modification entries were further extended using the G-PTM-D strategy in bottom-up proteomic analysis.

**Figure 2:**
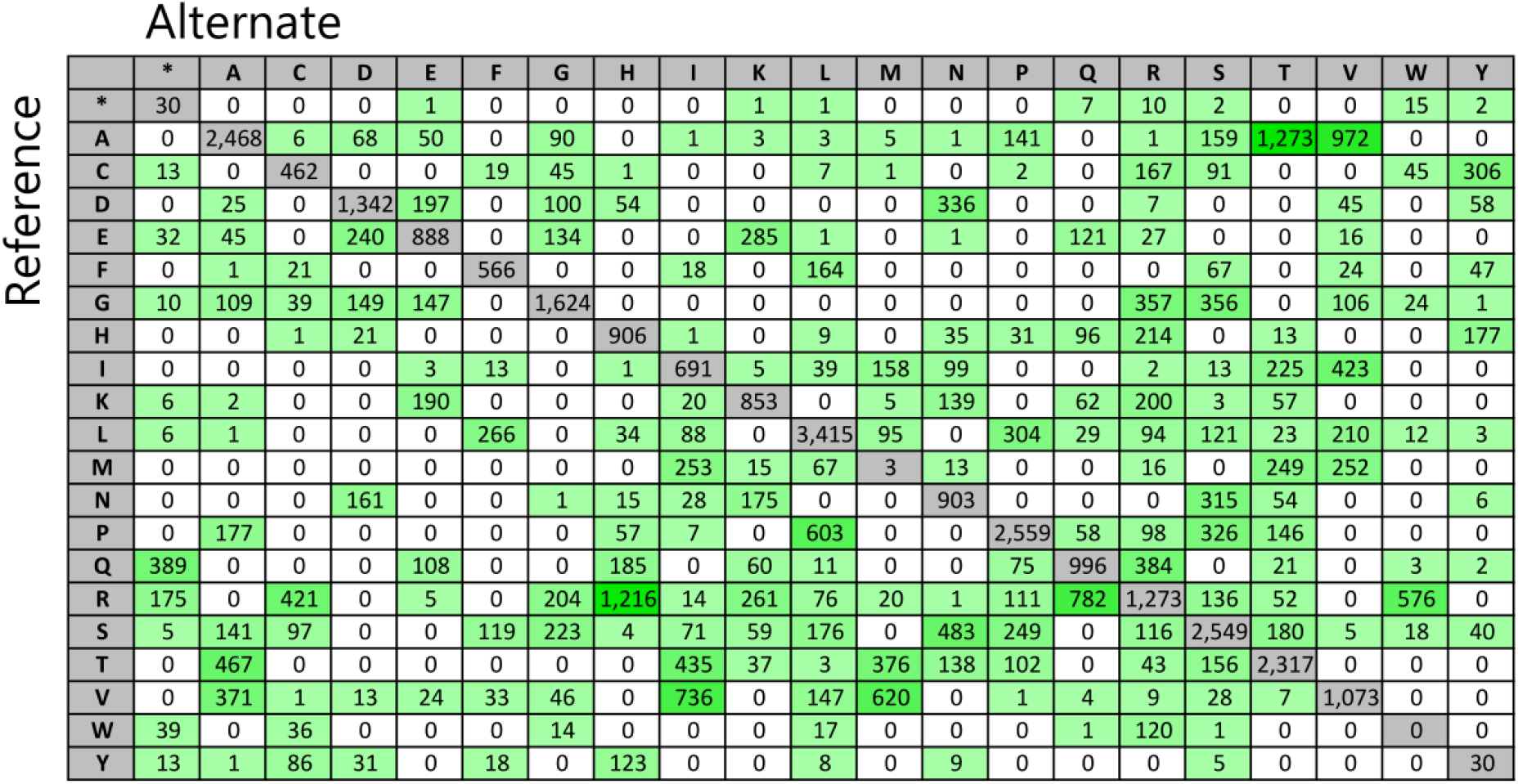
(Two columns) Coding variations were applied to proteins generated from the Ensembl reference genome including the amino acid substitutions listed here (green), while synonymous variations were excluded (grey).

**Figure 3:**
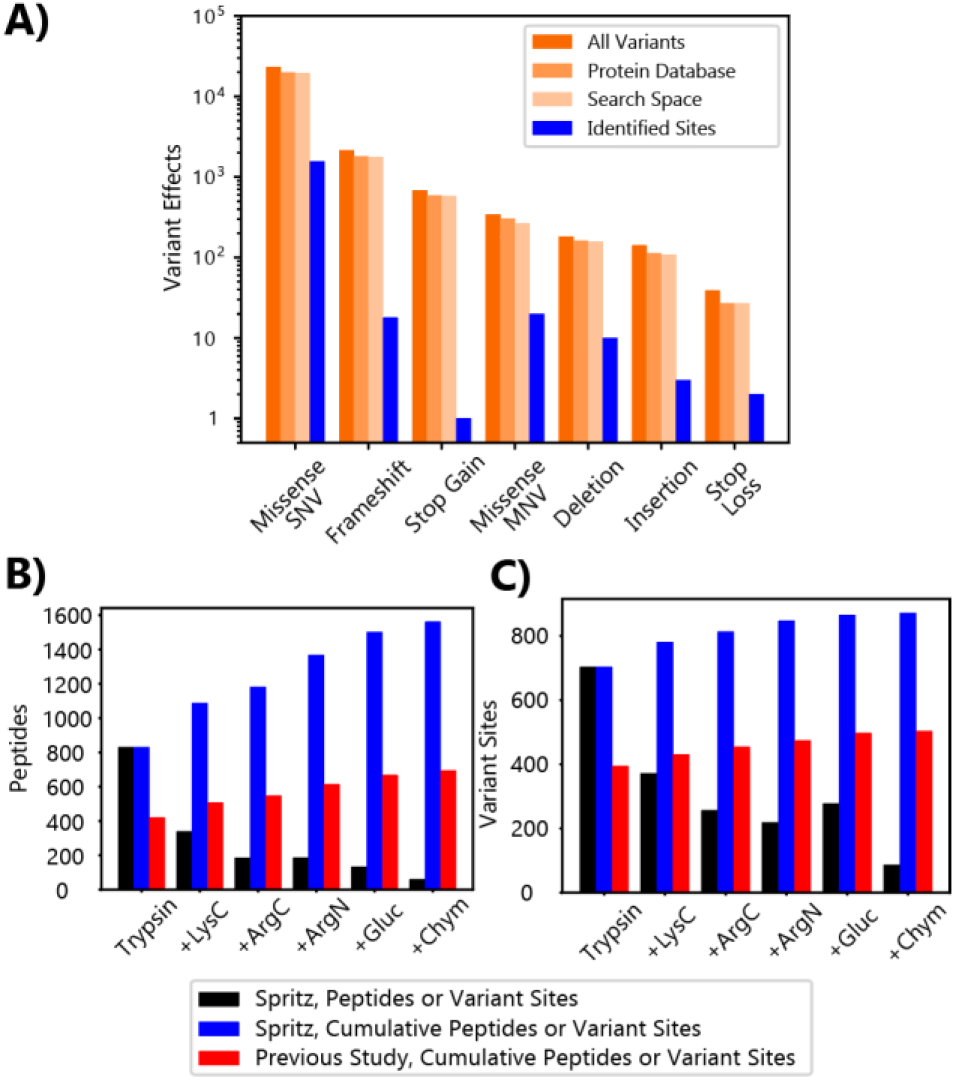
(One column.) A) Several types of variants were called by GATK, annotated in the protein database, included in the search space, and identified by the bottom-up proteomic analysis of Jurkat lysate. B) Comparison of the number of variant peptides and C) variant sites identified by Spritz analysis (blue) and by the previous analysis ^11^ (red). The uniquely identified variant peptides or sites for each protease are shown in black, whereas the cumulative total of variant peptides or sites are shown in blue or red. As reported previously ^35,36^, the use of multiple proteases enhances sequence coverage, leading to additional variant identifications, as shown in B and C.

### Variant Peptide Identifications

The Spritz-generated proteogenomic databases facilitate comprehensive detection of variant peptides by including many types of sequence variations including often neglected indels and frameshifts. A total of 196,056 peptides were identified (representing a 0.5% increase over using only protein sequences generated from the reference genome), including 1,578 variant peptides identified at 1% global FDR (Table S3-S4). These peptides have a 1.4% group FDR indicating that variant peptide identification has similarly good false positive rates to peptide identification overall. The majority of sequence variations were also validated by the fragmentation data; 98% of sequence variations have peptide fragment evidence, and 60% are validated by the stricter measure of sequential fragment ions covering the sequence variation. The total number of variant peptides (1,578) is over two times as large as the number of variant peptides identified in the previous analysis^11^ of these data (695; Figure 3B). These peptides identified 872 unique genomic variant sites (*e.g.*, missense SNV, indel), nearly double the number found previously (Figure 3C). However, these sites represent only 9.4% of the 9,326 unique variant sites annotated by Spritz, a slight improvement over the 6.9% (395/5755) of nsSNVs detected previously. The increase in variant peptides and sites identified indicates that Spritz was successful in improving the search for variant peptides by utilizing updated computational genomic tools like GATK, including additional types of variations besides only missense SNVs, and using the search engine MetaMorpheus to identify modified variant peptides.

### Modified Variant Peptide Identifications

The combination of a Spritz database and the use of G-PTM-D within MetaMorpheus allows for identification of post-translationally modified variant peptides with the modification anywhere on the peptide, including the variant site. In addition to the 49,999 modifications of 114 types transferred from UniProt by Spritz, the G-PTM-D strategy ^17,20^ was used to annotate 376,399 modifications of 60 types, including 268 modifications annotated on variant sites. This database was searched with a narrow mass tolerance to identify 326 modified variant peptides (1.8% group FDR; Figure 4A; Table S5), 14 of which were modified on the sequence variation (Table S6).

**Figure 4.**
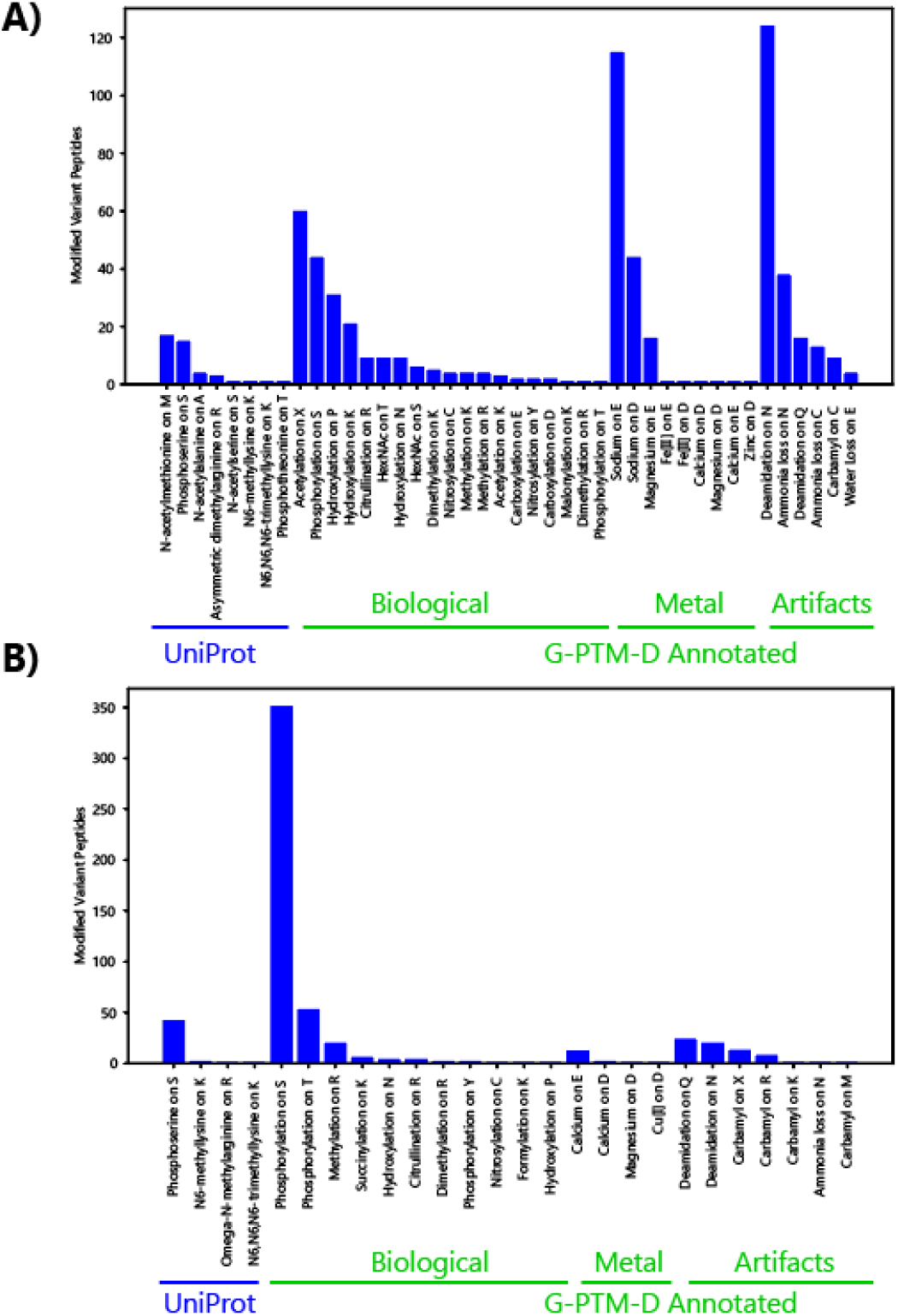
(One column.) Modifications detected on variant peptides in A) Jurkat multi-protease data and B) U2OS phosphoproteomic data.

Modified variant peptide identifications improved the overall detection of genomic variant sites. Of the 872 genomic variant sites identified by variant peptides, 7.1% (62 sites) were identified by modified variant peptides only.

To investigate modified variant peptides in samples enriched for a modification, we analyzed a U2OS RNA-Seq^22^ and phosphoproteomic dataset^21^. The RNA-Seq dataset had 10,247,413 reads; after trimming and filtering, 10,0253,77 reads remained, the mean per base quality score increased from 15.2 to 16.4, and 95.7% of reads were aligned to the genome by *hisat2.* Spritz analyzed these alignments to provide 56,072 variant calls. Despite being a smaller dataset than used for Jurkat (8.8% read count, 15.1% variant calls, 20.0% PSMs, 26.1% peptide IDs, trypsin only), a similar number of modification candidates were localized to the variant sites. A total of 241,027 modifications were added to the database, including 209 modifications annotated on variant sites. These candidate modifications on variant sites were 69.3% phosphorylations compared to 11.9% for the bulk Jurkat experiment, as expected for a phosphoproteomic dataset. Searching this annotated database yielded 248 modified variant peptide identifications (0% group FDR, *i.e.,* no decoys of this type were found at 1% FDR; Figure 4B; Table S7-S9), including 25 with phosphorylated variant sites (Table S10). This suggests that 11.8 times more modified variant sites are found in phospho-enriched samples relative to the total variants called for the sample. One example is the identification of a phosphoserine at residue 441 of Pinin, the site of a T > S variation, near other known phosphosites^37^.

### Proteoform Identification with Sequence Variations

Spritz facilitates top-down proteoform analysis by streamlining variant detection from RNA-Seq data and by running custom scripts to generate a protein database listing full-length protein sequences with annotations of variants and modifications. We used a new top-down search in MetaMorpheus to search 11 GelFrEE fractions of Jurkat cell lysate using a Spritz database. This “pruned” database was annotated with modifications using G-PTM-D with bottom-up data and pruned for modifications detected at 1% FDR. (We note that using the “protein pruned” database, which has only proteins detected at 1% FDR in bottom-up data, led to 15% fewer proteoform identifications.) This analysis resulted in the identification of 1,392 proteoforms including 30 variant proteoforms at 1% global FDR (0% group FDR) and 43 variant proteoforms at 1% group FDR (Table S11-S12). These 43 proteoforms originate from 13 proteoform families corresponding to 13 genes. Two examples are provided in Figure 5, including the SEC61B proteoform family where we observed both alleles of a heterozygous variation (Figure 5A) and the H3F3C proteoform family, where many proteoforms were detected from a sequence with two homozygous variations (Figure 5B).

**Figure 5.**
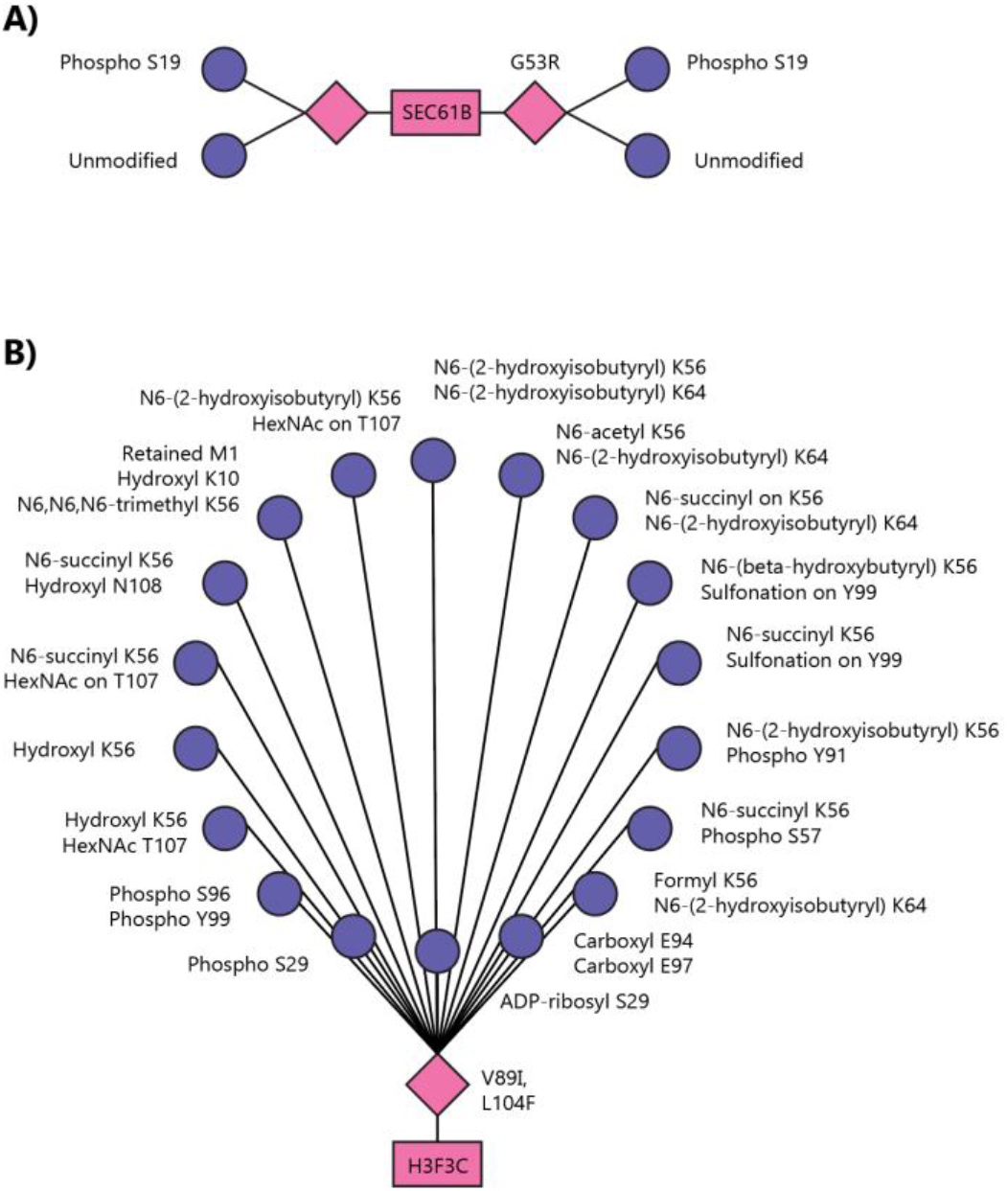
(One column) Manually constructed proteoform family examples containing variant proteoforms. Each pink square represents a gene, each pink diamond represents a unique transcript (e.g., variant, isoform), and each purple circle represents a proteoform identification. A) Proteoform family containing proteoforms generated by translation of the reference transcript as well as the variant transcript. All of these proteoforms are still related to a single gene and therefore belong to the same proteoform family. B) Proteoform family for a histone containing two single amino acid variants, and multiple post-translational modifications.

## Discussion

We report a new tool named Spritz that creates proteogenomic databases for the identification of variant peptides and proteoforms. Using a deep RNA-seq dataset for Jurkat cells, we constructed a proteomic database annotated with the coding effects of missense SNVs, missense MNVs, indels, frameshifts, stop loss, and stop gain variations. We demonstrate the improved capacity of Spritz-constructed databases to detect variant peptides using deep bottom-up proteomic data and the identification of variant proteoforms in top-down proteomic data for Jurkat cell lysate.

In particular, we show how modified variant peptides detected in unenriched samples improve the detection of variant sites by around 7%. This improvement was enabled by the application of the G-PTM-D search strategy within the search engine MetaMorpheus, as well as the customized UniProt-XML format reported in this work that enables annotating modifications on amino acids within the site of variation.

To further explore the detection of modified variant sites, we analyzed a phosphoproteomic dataset for U2OS. We found 11.8 times more modified variant sites than in the unenriched Jurkat dataset. This indicates the construction of proteogenomic databases, particularly from deep RNA-Seq or exome sequencing datasets, may be important for comprehensive detection of phosphosites.

One of the important motivators for developing Spritz was the need to accurately construct full-length variant protein entries to enable the detection of variant proteoforms. We used an extended version of the variant annotation program SnpEff to output full-length proteins annotated with variants in the customized UniProt-XML format, and we combinatorially add these variants to the protein sequences within MetaMorpheus. This enabled us to find 43 variant proteoforms, including proteoform families like that of H3F3C, which would not have been detected without including homozygous sequence variations.

Finally, Spritz was constructed with user friendliness in mind in an effort to make proteogenomic databases more accessible to the proteomics community. Because the proteomics community is the target audience for this tool, we constructed a GUI for running this tool in Windows, built upon a Docker container. The RNA-Seq data files and settings chosen in this tool are output into a config file that directs the downloading of reference files, as well as the setup and execution of approximately 20 tools to construct a sample-specific proteogenomic database. We hope this eases the learning curve for utilizing proteogenomic strategies and allows the proteomic community to more easily ask questions requiring proteogenomic workflows.

## Abbreviations

GUI: graphical user interface
PTM: post-translational modification
VCF: variant call format
BAM: binary alignment map
SAAV: single amino acid variation
SNV: single nucleotide variation
MNV: multi-nucleotide variation
SRA: sequence read archive
RPLC: reverse phase liquid chromatography
gVCF: genome variant call file
G-PTM-D search strategy: global post-translational modification discovery search strategy
FDR: false discovery rate
IDs: identifications
GATK: genome analysis toolkit

## Acknowledgments

Research in this publication was supported by the National Cancer Institute (NCI) of the National Institutes of Health (NIH) under award number R01CA193481. A.J.C. acknowledges support of the Computation and Informatics in Biology and Medicine training grant from National Library of Medicine (NLM) of the NIH, award number 5T15LM007359. R.M.M. was supported in part by the NIH Chemistry–Biology Interface Training Grant, T32 GM008505. The content is solely the responsibility of the authors and does not necessarily represent the official views of the NIH. A.J.C. acknowledges continued support from Emma Lundberg and funding by the Knut and Alice Wallenberg Foundation (2016.0204) and Swedish Research Council (2017-05327). We also acknowledge the other members of the Smith Lab software team, Zach Rolfs and Leah V. Schaffer, for contributing daily to maintaining the software used in this project.

## Author Contributions

AJC conceived of and wrote Spritz. AJC adapted the UniProt-XML format to accommodate modified variant annotations. AJC and RMM adapted MetaMorpheus to enable the G-PTM-D strategy to discover modifications of variant protein sequences. AJC and RMM developed scripts to count the number of variations detected by MetaMorpheus and debugged the integration of MetaMorpheus and Spritz. AJC ran Spritz to generate the proteogenomic databases. AJC and RMM analyzed the bottom-up and top-down proteomics datasets with Spritz databases. KI and LL created the Spritz GUI. KI extended the *snakemake* workflows for increased ease of use. RJM developed top-down MetaMorpheus analysis. AJC and RMM drafted the manuscript. MRS, BLF, and LMS advised on the project and coordinated funding. All authors edited and approved the submitted manuscript.

## Competing Financial Interests

The authors declare no competing financial interests.

## Supporting Information

Supporting Tables (XLSX Format):

- Table S1. Data sets.
- Table S2. Search settings.
- Table S3. Variant Peptides Detected in Bottom-Up Jurkat Data.
- Table S4. Variant Sites Detected in Bottom-Up Jurkat Data.
- Table S5. Modified Variant Peptides Detected in Jurkat Bottom-Up Data.
- Table S6. Modified Variant Sites Detected in Bottom-Up Jurkat Data.
- Table S7. Variant Peptides Detected in U2OS Phosphoproteomic Data.
- Table S8. Variant Sites Detected in U2OS Phosphoproteomic Data.
- Table S9. Modified Variant Peptides Detected in U2OS Phosphoproteomic Data.
- Table S10. Modified Variant Sites Detected in U2OS Phosphoproteomic Data.
- Table S11. Variant Peptides Detected in Top-Down Jurkat Data.
- Table S12. Variant Sites Detected in Top-Down Jurkat Data.

## Table of Contents Graphic

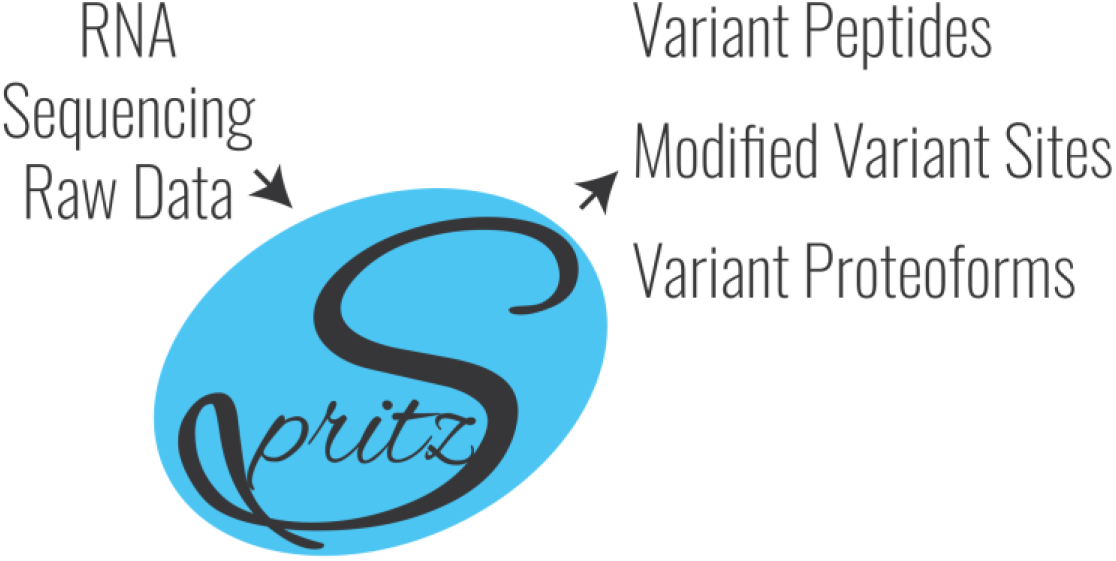

## Notes

### Competing Interest Statement

The authors have declared no competing interest.

